# Learning to avoid collisions: Bees use cues perceived before and after a collision to prevent negative consequences during navigation in cluttered areas

**DOI:** 10.1101/2025.08.27.672532

**Authors:** Manon Jeschke, Maximilian Stahlsmeier, Olivier J. N. Bertrand, Martin Egelhaaf

## Abstract

Bumblebees navigate complex environments where collisions with obstacles can impair flight performance. While bees possess innate collision avoidance reflexes, they may benefit from learning to identify and avoid high-risk collision areas using environmental cues. However, the temporal dynamics of how bees form associations between visual cues and collision experiences remain unclear. We investigated whether bumblebees associate visual cues perceived before or after collision events with a movement direction. Individual foragers were trained to navigate through a flight tunnel containing a transparent barrier and an LED panel that switched colours immediately after the bee’s first collision. Following training, we tested bees’ responses to each colour cue without the barrier. Bees demonstrated significant preferences for avoiding the previously blocked side when presented with either the pre-collision colour (81% correct responses) or post-collision colour (71% correct responses), while showing no preference when no colour cue was presented. Individual analysis revealed that 61% of bees responded to both cue types, while others showed selective responses to specific temporal windows. These results demonstrate that bees can form associations with visual cues encountered in different time windows relative to negative experiences, revealing temporal flexibility in associative learning that contributes to successful navigation in cluttered environments.

## Introduction

Bumblebees navigate through complex environments where the risk of collisions with obstacles is omnipresent. These collisions can have serious consequences, potentially causing injuries or wing damage that impair flight performance (Foster & Cartar, 2011; Mountcastle et al., 2016). Despite these challenges, bumblebees demonstrate remarkable navigational abilities, successfully traversing cluttered environments such as dense vegetation, urban landscapes, and artificial structures to reach foraging sites and return to their colonies (Burnett et al., 2021; Crall et al., 2015; Ravi et al., 2022; Woodgate et al., 2016).

In addition to their navigational capabilities, bees demonstrate excellent learning abilities, readily forming associations between rewards and diverse environmental cues (Avarguès-Weber et al., 2011; Giurfa & Sandoz, 2012; Laska et al., 1999; MaBouDi et al., 2025). This associative learning enables them to identify, memorize, and discriminate rewarding outcomes, including high-quality food sources and safe routes between food locations (Buatois & Lihoreau, 2016; Lihoreau et al., 2010, 2012).

Recent studies have revealed sophisticated learning mechanisms that enable bees to navigate safely through cluttered environments and establish habitual routes, even if the obstacles were transparent and only barely visible and were thus unable to trigger any visual collision avoidance mechanism (Bertrand et al., 2021; Gonsek et al., 2021; Jeschke et al., 2025). This adaptive capacity allows bees to minimize collision risks while maintaining efficient foraging behaviour. Thus, it was shown that bees can associate colour cues with specific movement directions to effectively navigate through tunnels partially obstructed by transparent barriers (Bertrand et al., 2021). However, this study left open the question of at what time window during the collision experience bees form these critical associations between visual cues and avoidance behaviour.

Collision avoidance in bees is primarily controlled by innate reflexive behaviours that use optic flow information to assess object proximity and navigate around obstacles (Egelhaaf, 2023; Ravi et al., 2022). These visual collision avoidance systems work effectively, even enabling successful navigation through highly cluttered environments (Bertrand et al., 2015; Burnett et al., 2021; Jeschke et al., 2025). However, real-time collision avoidance is costly for the visual system. Bees must continuously assess gap widths between obstacles using motion parallax information (Ravi et al., 2019), requiring constant distance estimation and gap assessment in complex environments. Since collision failures can have serious consequences, additional learning mechanisms that help bees avoid high-risk collision areas in the environment may be beneficial. Such learned associations between high-risk areas (e.g., dense vegetation patches like bushes on a meadow) and visual cues (e.g., colour of near-by flowers or prominent landmarks) could allow bees to plan routes and stay away from these high-risk areas during their flight.

The timing of associative learning in collision avoidance has important implications for understanding how bees process and respond to environmental challenges. In classical conditioning, the conditioned stimulus (e.g. a colour cue) precedes the unconditioned stimulus (e.g. a food reward) and may even overlap with it in time (Akpan, 2020). However, bees are also capable of forming associations under more complex temporal conditions. They can learn when there is no temporal overlap between the conditioned and unconditioned stimulus (“trace learning”) (Szyszka et al., 2011; Wystrach et al., 2020), when the conditioned stimulus follows the unconditioned stimulus (“backward learning”) (Hussaini et al., 2007) or even in the absence of explicit reinforcement (“latent learning”) (Wystrach, 2023). In foraging contexts, where bees learn to associate colour or spatial cues, learning critically depends on whether the stimulus is perceived during arrival or during the departure phases of their visits to a rewarding place (Lehrer, 1993, 1994). These experiments involved positive reinforcement learning, where colour cues were directly associated with a reward. However, learning to avoid collisions with barely visible objects presents a fundamentally different ecological context: if bees want to avoid collisions with such objects, they can do so not through positive reinforcement, but through negative reinforcement learning, in which the primary feedback comes from the adverse consequences of collisions with obstacles.

Evidence shows that insects can learn from negative experiences to modify their future behaviour (Chittka et al., 2003; Avarguès-Weber et al., 2010; de Brito Sanchez et al., 2015; Nityananda & Chittka, 2015). Ants that fall into invisible traps during homeward journeys reshape their routes and avoid traps in subsequent trials, showing increased scanning behaviour in areas preceding previously encountered traps, suggesting that visual scenes experienced before negative events act as aversive feedback (Knaden, 2020; Wystrach et al., 2020). Similarly, when bees collide with transparent obstacles, they could potentially form associations with salient objects during two time windows: before the collision (linking pre-collision visual cues with the negative experience) or after the collision (when scrutinizing the environment to identify solutions and associate post-collision cues). However, because visual environments remain constant in both cases, the exact temporal window in which these insects form negative associations remains speculative.

To address this problem, we designed an experimental paradigm that allows us to distinguish between associative learning before or after the first collision with an obstacle. We trained bees to navigate through a flight tunnel containing an LED panel capable of displaying different colours and a transparent barrier that partially blocked their path. Critically, the LED panel displayed one colour before a bee’s first collision with the barrier and immediately switched to a different colour after the first collision occurred. This design enabled us to test whether bees associate visual cues perceived before the first collision with the collision experience, or whether they form associations with cues encountered after the first collision. Therefore, after successful training, we removed the transparent barrier and repositioned the LED panel to assess which temporal association had been formed. By presenting either the pre-collision or post-collision colour and observing the bees’ subsequent flight paths, we could determine whether bees had learned to associate the visual cues perceived before and/or after collision events with appropriate directional decisions for collision avoidance.

## Methods

### Animal Handling

Two commercial bumblebee colonies (*Bombus terrestris*) were obtained from Koppert B.V., The Netherlands. Upon arrival, colonies were transferred into custom-designed acrylic housing boxes (24 × 24 × 40 cm^3^) that were draped with black cloth to maintain darkness and simulate natural nesting conditions. Within these enclosures, bees were provided with pollen *ad libitum* and had access to 30% sucrose solution in a designated foraging area.

Foraging bees that demonstrated consistent flight activity between the hive and the feeding chamber were selected for individual marking. The marking procedure involved capturing individual bees and temporarily immobilizing them within a marking tube. Numbered coloured plastic tags were then affixed to each bee’s thorax using resin to enable individual identification throughout the experimental period. Following the marking procedure, bees were released back into the experimental setup at the nest entrance.

### Experimental Setup

The experimental setup consisted of two flight tunnels connecting the bumblebee hive to a foraging chamber (Figure 1). Bees accessed the flight tunnels from the hive through transparent tubes (2.5 cm diameter) and small acrylic transfer boxes (8 × 8 × 8 cm^3^). Each flight tunnel measured 200 × 50 × 30 cm^3^, with walls and floor covered with a red and white 1/f noise pattern as described by Ravi et al. (Ravi et al., 2019) to provide bees with a naturalistic spatial frequency spectrum (Schwegmann et al., 2014). The tunnel ceiling was made of transparent acrylic to facilitate behavioural observation.

**Figure 1.**
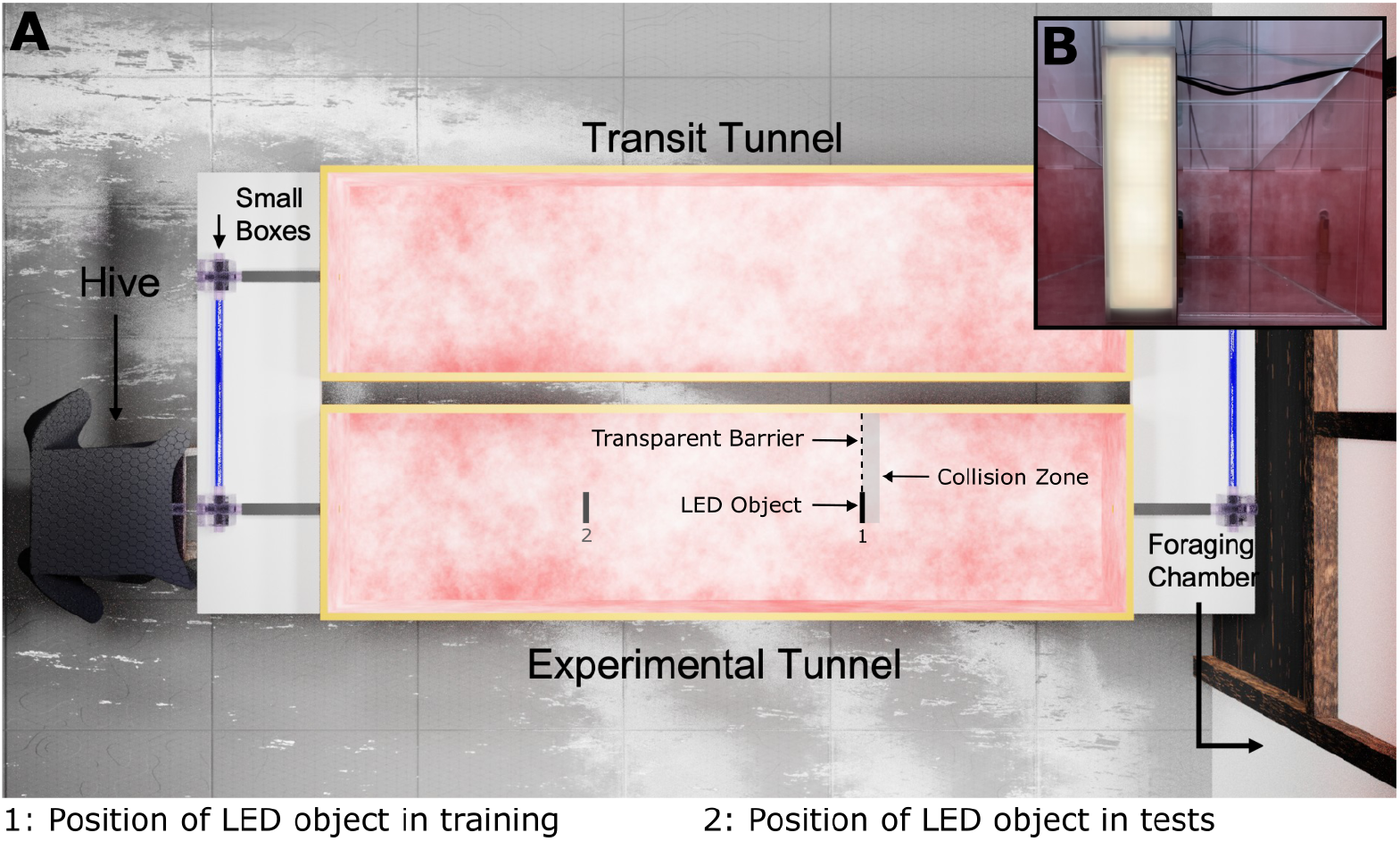
Experimental setup. A: Rendered top view of the setup that was recreated using the graphics software *Blender*. The hive was connected to the setup via small transparent boxes and tubes. Bees entered the foraging chamber (not shown) by flying through the transit tunnel (top). For the experiments, returning foragers were redirected into the experimental tunnel (bottom), which they had to cross to return to their hive. During training, the experimental tunnel was equipped with an LED panel at position 1 that could display different colours and a transparent barrier which blocked half of the tunnel. Once a bee entered the collision zone and collided with the barrier or the object for the first time, the colour of the LED object changed. For the tests, the LED panel was displaced to the test position at position 2 and the transparent barrier was removed. B: Photographs taken from within the tunnel during the training with the LED panel in the centre and the transparent barrier to the left.

Bees that left the hive flew through the empty transit tunnel into the foraging chamber, where they could feed on sugar solution provided in gravity feeders. Upon returning from the foraging chamber, individual foragers were redirected into the experimental tunnel which they had to cross to return to their hive. The experimental tunnel contained two key components: an LED panel (8 cm × 30 cm × 1.5 cm) and a removable transparent acrylic barrier (21 cm × 30 cm × 0.3 cm) that could block half of the tunnel width. The LED panel consisted of four LED matrix panels with 64 LED lights each, covered by an acrylic milk glass plate. The LED panel was controlled by an Arduino microcontroller and capable of displaying different colours as specified by the experimental protocol. The spectral reflectance of the colour cues used in the experiment was quantified with a spectrometer (OpticOcean4000; see Suppl. Figure 1). Flight behaviour was captured using four synchronized high-speed cameras (Basler acA2040-90umNIR) equipped with red filters (Heliopan RG715). Two cameras provided top-down views from 2 m height, each covering half the tunnel with 25 cm overlap. Two additional cameras were positioned at the opposite ends of the experimental tunnel at 1.8 m height and angled at 45°, providing longitudinal side views with central overlap. Camera calibration was performed to determine intrinsic parameters (focal length, optical centre, and lens distortions) and extrinsic parameters (camera orientations and positions) using standard checkerboard calibration methods implemented in OpenCV software.

The room and the flight tunnels were illuminated by daylight. To enhance bee visibility in video recordings, the tunnel received additional illumination from below through a 650 nm cutoff low-pass acrylic filter (following (Eckel et al., 2023)). This configuration created a bright background for camera detection while remaining imperceptible to bees due to their limited sensitivity in this wavelength range (Skorupski & Chittka, 2010), thereby maximizing contrast for bee detection. Images were acquired at 100 frames per second, enabling detection, tracking, and three-dimensional trajectory reconstruction of individual bees.

### Experimental Procedure

Prior to the experiment, the bees were given a one-week habituation period to acclimate to the experimental setup, allowing them to freely navigate in the transit tunnel and enter the foraging chamber to forage for their colony. Afterwards, trials were conducted using forager bees that demonstrated consistent travel between the hive and foraging chamber. The experimental protocol consisted of three phases: pre-training, training, and testing.

During pre-training, the LED panel was positioned at the training location (Figure 1, Position 1) but remained switched off. Individual bees completed three pre-training trials in which they traversed the experimental tunnel, starting at the foraging chamber, to return to their hive. We recorded their flight trajectories and determined on which side of the LED object each bee chose to pass, allowing assessment of any inherent population-level side bias.

The training phase consisted of five trials per bee with the LED object activated and the transparent barrier positioned to block half the tunnel width. The barrier position was randomly assigned for each individual bee to either the right or the left of the LED display but remained unchanged for that bee throughout the training period to allow the bee to associate the colours of the LED panel with the side of the obstacle. The colour of the LED display panel could alternate between yellow (hex-code #d49e2a) and purple (hex-code #9e2ad4) (see spectrometer measurements in Suppl. Figure 1), with one colour designated as the “before” colour and the other as the “after” colour. Note that half of the bees were trained with yellow as the colour shown before a collision and purple as the colour shown after a collision and the second half of bees vice versa to ensure that successful training was not caused by a potential colour preference. When a bee entered the collision zone (defined as 2 cm before the transparent barrier) (Figure 1) and thus collided with the barrier for the first time, the experimenter immediately changed the LED colour from the before colour to the after colour. The transparent barrier was regularly cleaned with ethanol and dried to remove dust and ensure that bees could not detect the transparent barrier visually.

Following training completion, the LED panel was repositioned to the test location to exclude that bees associated external cues with the movement direction (Fig.1, Position 2). During test trials, the transparent barrier was removed, and the bees were presented with either the before or after colour from the LED panel, with the colour remaining constant throughout each trial. Control trials were conducted with the LED panel switched off to determine whether bees had learned to associate specific colours with directional choices or simply with the presence of the LED panel itself.

### Trajectory Recording and Reconstruction

The bees’ flight paths were recorded using custom-written C++ software (available at https://gitlab.ub.uni-bielefeld.de/neurobio-public/mcam-suite). The tracking procedure consisted of multiple sequential steps. Initially, a background reference image was established by averaging the first 100 frames of each recording. Real-time background subtraction was then implemented during recording to detect bee movement, generating cropped images containing only the differences between current and background frames together with the corresponding spatial coordinates. Image classification was subsequently applied to retain only frames containing bees. Three-dimensional flight trajectories were reconstructed through triangulation when multiple synchronized cameras simultaneously detected the same bee, enabling calculation of precise spatial positions within the tunnel environment. Complete documentation, including recording software code, camera calibration protocols, and trajectory reconstruction algorithms, is publicly available in our repository (https://gitlab.ub.uni-bielefeld.de/m.jeschke/learning_to_avoid_collisions). Post-processing involved filtering trajectories using a second-order low-pass filter with a 2 Hz cutoff frequency to remove high-frequency noise. We removed trajectories of bees from the analysis that dropped out of the experiment before completing the experimental protocol due to death, or those of bees that did not leave the hive or walked through the tunnel. Overall, we analysed 251 flights of 24 bees from three different hives.

### Data analysis and statistics

After reconstructing the trajectories, we could determine on which side the bees passed the LED object. For each flight we calculated the median y-position of the bees from 2.5cm before to 2.5cm after the LED object. If a bee flew back and forth and passed the LED object multiple times, we only considered the first approach in our analysis. After verifying that the behaviour of the different bees was consistent regardless of whether the transparent barrier was placed on the left or right side of the LED panel during the training phase for the respective bee, we pooled the data and mirrored the trajectories of the bees that were trained to pass the LED panel on the right side for analysis. The bees could either pass the LED panel on the side that was ‘blocked’ during the training (i.e. where the transparent barrier was placed) or on the ‘clear side’ (i.e. where no barrier was placed during the training). We analysed the trajectories and determined the side choices for all flights in the pre-training, training and tests. We tested with a *X*^2^ − *test* if there was a preference for either the blocked or clear side in the tests (‘left’ and ‘right’ in pre-training) if they were free to choose.

Furthermore, we analysed the side chosen by the bees in the training phase and calculated the proportion of correct choices, i.e. instances where the bees passed on the clear side without colliding with the transparent barrier during their initial approach. This was calculated for each training trial. We also calculated how long it took the bees to complete the training trial and to cross the tunnel as well as the straightness of the paths. The path straightness is defined as the ratio of the shortest distance between tunnel entrance and exit divided by the actual flown distance by the bee, resulting in a value between 0 and 1. To describe the relationship between the individual flight characteristics and the number of consecutive trials, and thus the increasing training experience, a linear regression and a Spearman correlation coefficient was calculated. To test whether flight characteristics differ between the test conditions while the bees approached the LED panel, we calculated the time needed to cross the tunnel and the path straightness for each test flight and performed a Kruskal-Wallis-test.

## Results

### Bees associate cues they perceive before and after the first collision with a movement direction

To test whether bumblebees associate visual cues perceived before or after a collision with the negative experience of colliding with a transparent obstacle - and whether this association enables them to avoid risky areas for collisions in the future - we trained bees in a flight tunnel containing a colour-changing LED panel and a transparent barrier. The bees had to navigate around the barrier to cross the tunnel and return back to their hive. Prior to the training, each bee undertook three pre-training flights, during which the tunnel contained only the turned-off LED panel. This was to assess if the bees, as a population, had a side bias or passed the LED panel evenly on both sides. Bees showed no statistically significant directional preference when navigating past the LED panel, with 41% of choices to pass on the left and 58% of choices on the right (*X*^2^ − *test*, t = 1.724, p = 0.189, Figure 2A). This baseline established that the population of tested bees did not show a systematic bias before experiencing the experimental conditions.

**Figure 2.**
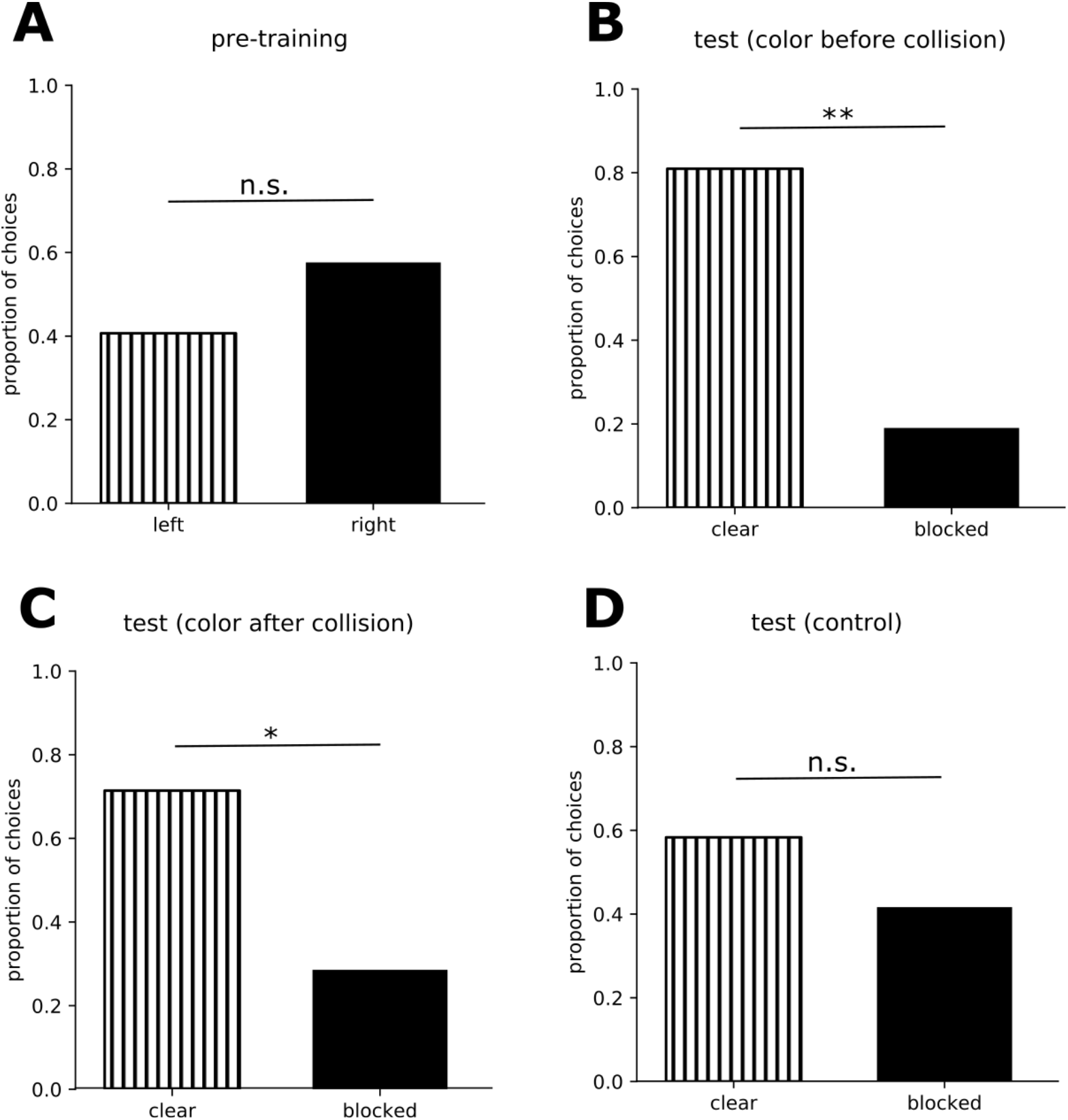
Choices of bees in the tests and pre-training. The bar plots show on which side the bees choose to pass the LED panel. The LED panel displayed either the colour shown before a collision during training (B) (n=21), the colour that was shown after a collision during training (C) (n=21), or the LEDs were turned off (D) (n=24). The bees could choose to pass the LED panel either on the side that was blocked by a transparent barrier during the training or on the clear side which was not blocked during the training. In the pre-training, the bees did not experience a barrier yet and freely chose to pass the turned off LED panel on the left or right side (A) (n=59). A *X*^2^ − *test* was used to test whether the distribution of choices is different from chance level.

During the tests, bees’ side choices varied depending on the LED cue presented. When the LED panel displayed the colour that had been shown before collisions during training (pre-collision cue), bees demonstrated a strong preference for passing on the clear side (81% of choices, *X*^2^ − *test*, t = 8.047, p = 0.004, Figure 2B). When presented with the colour that appeared after collisions during training (post-collision cue), the bees also showed a preference for the clear side, even if this was somewhat weaker (71% of choices, *X*^2^ − *test*, t = 3.857, p = 0.049, Figure 2C). To control whether the bees associated the colour of the LED panel with the avoidance maneuver and not the object itself, we turned the LED panel off and provided no colour cue in the control. In this condition bees again showed no statistically significant side preference (58% of choices for the clear side vs. 41% of choices for the blocked side, *X*^2^ − *test*, t = 0.666, p = 0.414, Figure 2D).

When we analysed the side choices of individual bees, we found that 61% of all bees chose to pass on the clear side in both tests, while 17% passed on the clear side only when the pre-collision colour cue was presented in the test. 11% of bees passed on the clear side only when the post-collision colour cue was presented in the test. The remaining 11% of bees didn’t pass on the clear side of the tunnel, regardless of which colour cue was presented in the tests.

In summary, the results of the different test conditions show that most bees learned to associate both colour cues - the cues that were shown before the first collision and after the first collision - with the correct manoeuvre to avoid collisions with the transparent barrier; yet some bees appeared to use only either the pre-collision or the post-collision colour to avoid collisions in the tests.

### Bees improve their flight paths with increasing experience during training

Bees successfully learned to navigate the experimental setup in the training phase. The improvement in flight characteristics was visually apparent when comparing trajectories from early versus late training trials (Figure 3A). Initial flights were highly variable, and bees meandered with multiple course corrections and direction changes. We observed that the bees explored the area near the LED panel and transparent barrier and often collided with the latter. In contrast, final training flights were less convoluted and showed more direct trajectories with only occasional collisions with the transparent barrier.

**Figure 3.**
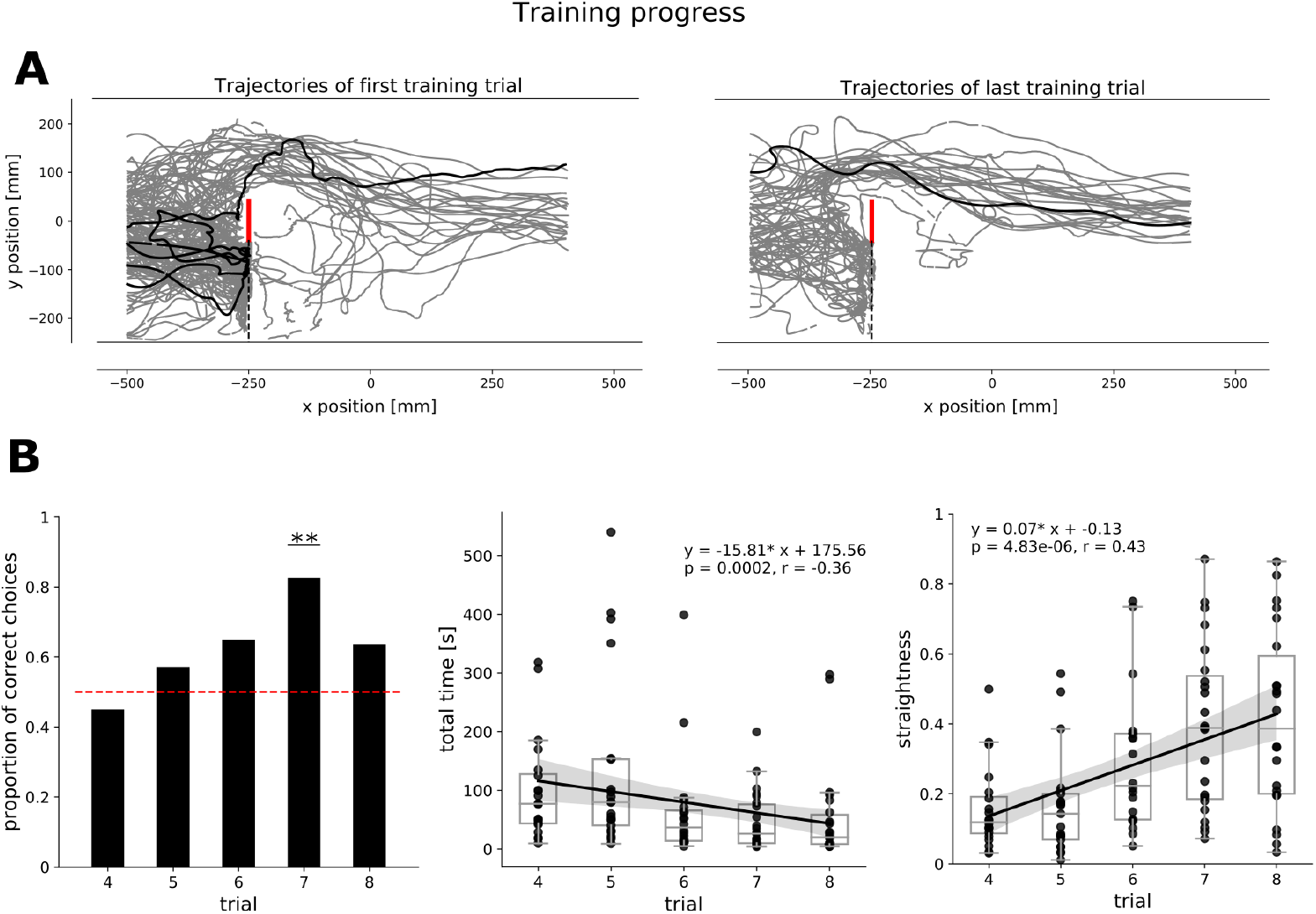
Behaviour of bees in the training phase. A: Trajectories of bees in the first and last training trial. Trajectories of bees in the section of the tunnel where the LED panel was placed are shown in grey. One example trajectory is highlighted in black. The bees flew from left to right. The LED panel is shown in red and the transparent barrier is indicated by a black dashed line. B: The barplot shows the proportion of correct choices in the training trials where the bees chose to pass the LED panel on the clear side during their first approach. The boxplots show the time bees need to cross the tunnel (n=107) and the straightness of the flight paths (n=107) with increasing experience in the training. A linear regression was fitted to the data, and a correlation was described using the Spearman correlation coefficient.

The proportion of correct choices, defined as passing the LED panel on the clear side during the first approach, increased progressively throughout training trials and reached up to 80% of correct choices (Figure 3B). After four training trials, the choice of the bees for the correct side differed significantly from chance level (*X*^2^ − *test*, t = 9.782, p = 0.0017). This learning dynamics demonstrated that the bees could associate visual cues with turning to avoid collisions.

By analysing several flight characteristics of training flights, the overall flight efficiency was found to improve with increasing experience. The time bees needed to cross the tunnel decreased on average by 74% from 76.92 seconds to 20.08 seconds (linear regression: y = -15.81x + 175.56, p = 0.0002, r = -0.36), while path straightness increased on average by 69% from 0.11 to 0.38 (linear regression: y = 0.07x - 0.13, p = 4.83 x 10^-6, r = 0.43). Thus, the bees became both faster and more direct as they gained more experience during the training.

### Approach behaviour of bees differs between tests

By analysing the bees’ approach behaviour during the tests, i.e. their trajectories from the tunnel entrance up to the point at which they reached the position of the LED panel, we revealed distinct patterns in both path straightness and approach time depending on the colour cue presented in the different conditions (Figure 4). Bees approaching the turned-off LED panel in the control condition showed the highest path straightness (median = 0.68), while those encountering the post-collision colour cue had the lowest path straightness (median = 0.55). In the pre-collision condition, bees displayed an intermediate path straightness (median = 0.58). Despite these numerical differences, the differences in path straightness were not statistically significant differences between the test conditions (Kruskal-Wallis test: t = 4.49, p = 0.1).

**Figure 4.**
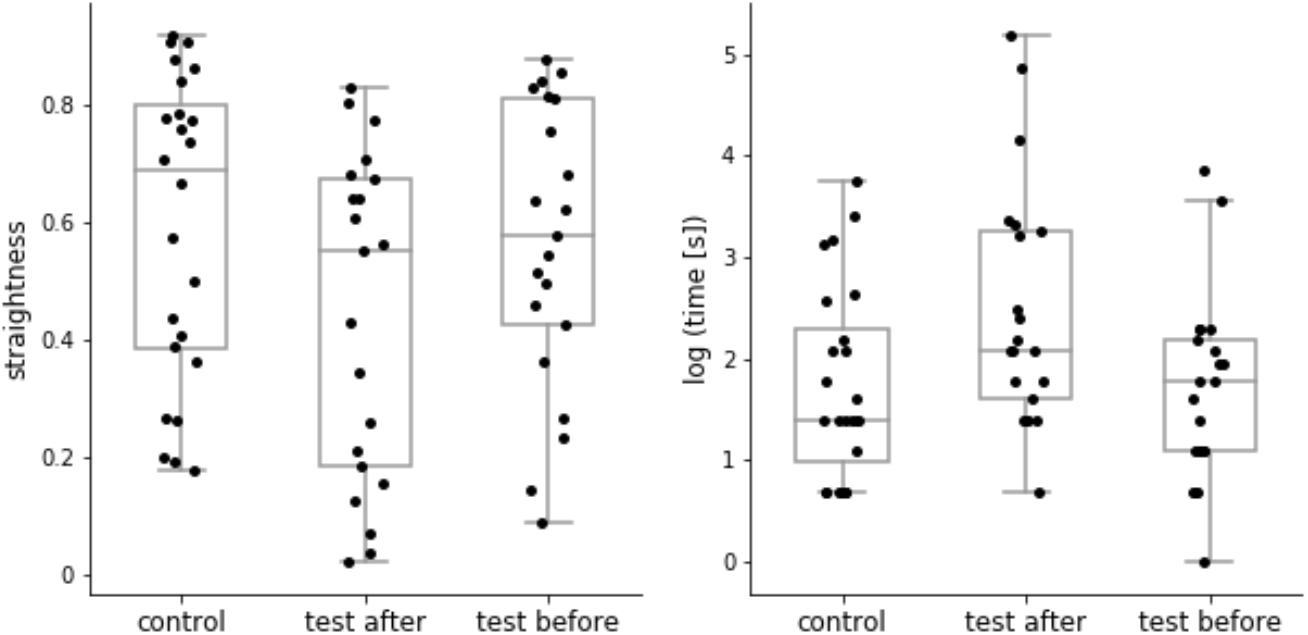
Flight characteristics of bees when they approach the LED panel in the test conditions. The boxplots show the path straightness (n=68) and the time (log-transformed) (n=68) the bees needed until they reached the LED panel in the test conditions. Differences between test conditions were tested using a Kruskal-Wallis-test.

The time it took the bees to approach the LED panel from the tunnel entrance also differed between conditions. The bees took the longest to reach the LED panel when the post-collision colour cue was presented (median = 8.47 seconds). When the pre-collision colour cue was displayed, bees approached the LED panel faster (median = 6.05 seconds), which is close to the time the bees needed in the control condition when no colour was displayed (median = 4.86 seconds). The Kruskal-Wallis test was used to compare the approach time in different conditions, revealing a trend for bees to take longer to approach the LED panel when the post-collision colour cue was presented, and to take longer to decide which side to pass the panel (Kuskal-Wallis test: t = 5.71, p = 0.057).

## Discussion

Our results demonstrate that bumblebees can successfully associate the unpleasant experience of colliding with transparent obstacles in a flight arena with specific colour cues in their vicinity. They then use this learned association during subsequent flights to avoid high-risk areas for collisions. Furthermore, during training the bees demonstrated a notable improvement in navigation efficiency. In doing so, the bees used the colour information they had perceived during training before the first collision (pre-collision colour), as well as the colour displayed after the first collision (post-collision colour). These findings show that bumblebees can form associative memories using visual cues perceived in different time windows relative to the punishing feedback (i.e. before and after collision events) to guide collision avoidance behaviour.

### Learning through negative reinforcement

Our study provides clear evidence that bees can learn to identify and avoid high-risk areas through negative reinforcement, demonstrating that - depending on the behavioural context - immediate positive reward may not be necessary for forming associations. Unlike typical foraging scenarios where bees associate floral characteristics with rewards, our experimental paradigm shows that negative experiences such as collisions can equally trigger associative learning processes that help bees navigate through dangerous areas.

Within the framework of classical conditioning, the collision event served as the unconditioned stimulus, while the colour cue from the LED panel functioned as the conditioned stimulus. However, because we studied natural flight behaviour in a semi-naturalistic setup, we observe high variability in the bees’ behaviour that extends beyond standard conditioning paradigms. This variability reflects the complex nature of learning during active navigation, where bees must integrate multiple sensory inputs while maintaining flight control.

Previous research has established that bees can learn under diverse temporal conditions, including trace learning (where stimuli are separated in time) and latent learning (without explicit reinforcement) (Szyszka et al., 2011; Wystrach, 2023; Wystrach et al., 2020). Our findings extend this understanding by demonstrating that learning to recognize and avoid high-risk areas can occur across different time windows during the collision experience itself. The neurobiological basis for this flexibility likely involves the mushroom bodies, known as brain regions critical for associative learning, memory storage, and retrieval in insects (Fiala & Kaun, 2024; Webb, 2024). These structures integrate sensory inputs with valence signals, enabling the formation of associations between environmental cues and positive or negative experiences. The mushroom bodies are essential for visual memory encoding during navigation, as demonstrated by studies showing that navigation becomes impossible when these regions are damaged (Buehlmann et al., 2023; Kamhi & Barron, 2020).

### Temporal flexibility in associative learning

A key finding of our study is that bees learned to associate both colour cues, the ones before and after their first collision within a trial, with appropriate area avoidance responses. Since we changed the colour of the LEDs after the first collision, we cannot completely separate pre-collision learning from genuine post-collision learning, because some bees experienced further collisions after the colour cue had been switched after the first collision. However, our results provide important insights into the temporal dynamics of learning to recognize high-risk areas.

The high success rate (81%) for avoiding collisions when presented with the pre-collision colour cue strongly suggests that visual information perceived immediately before the first collision plays a crucial role in learning. This finding indicates that bees can rapidly form associations between environmental cues and impending danger, even during their initial encounter with a hazard.

The substantial success rate (71%) for the post-collision colour cue presents a more complex scenario. Since the colour displayed after the first collision also preceded subsequent collision events during training, we cannot definitively determine whether associations formed through true post-collision processing or through subsequent pre-collision experiences. However, the effectiveness of both colour cues suggests that bees are able to associate multiple visual stimuli presented at different time points with the same behavioural outcome, in this case, recognizing and avoiding high-risk areas to prevent collisions.

This temporal flexibility in learning resembles findings from other bee studies, such as Lehrer’s work showing that bees can learn different visual features both during arrival and departure from objects (Lehrer, 1994). Our results extend this concept to learning about risky areas, demonstrating that temporal flexibility in associative learning extends beyond navigation around the nest entrance.

An important question for future research concerns which specific collision characteristics trigger learning most effectively. Learning may not occur uniformly across all collision events but could depend on factors such as impact intensity, contact duration, or the bee’s behavioural state during collision. Similar findings in ants support this hypothesis. Not all individuals learned alternative routes after falling into traps, with some employing different strategies, suggesting that more severe or prolonged negative experiences create stronger aversive memories (Knaden, 2020; Wystrach et al., 2020).

### Individual variation in learning strategies

Our analysis of individual bee responses revealed considerable variation in learning strategies. While 61% of bees demonstrated a successful association with both cue types, others showed selective learning for either pre-collision (17%) or post-collision (11%) cues. This variation between individual bees suggests that bees may employ different cognitive strategies for processing collision-related information, potentially reflecting differences in sensory processing capabilities, attention allocation, or memory formation mechanisms. Such variation is consistent with findings in other insect learning studies, where individual differences in learning performance are commonly observed (Klein et al., 2017). Bees display differences in inter-individual learning performance in different learning paradigms and sensory modalities, which remain consistent across learning tasks (Finke et al., 2023). However, the number of test trials per bee in our study was not large enough relative to allow for a conclusive assessment of inter-individual differences in learning strategies for specific bees. However, this issue is important for future research.

### Ecological relevance and Conclusion

The ability to form multiple, temporally distinct associations with identical behavioural outcomes represents a sophisticated form of learning that could provide significant adaptive advantages in natural environments. Real-world navigation scenarios often present complex situations where the same navigational solution may be signalled by different environmental cues encountered at various time windows during approach to or departure from objects. This learning flexibility also provides redundancy in memory formation, potentially improving navigation safety when individual cues become unreliable or unavailable. For instance, if specific landmark flowers fade or are removed, alternative visual cues learned during different phases of the collision experience could still guide appropriate avoidance behaviour.

In conclusion, our study reveals that bumblebees employ sophisticated learning mechanisms, allowing them to use visual cues encountered at different times to recognise and avoid areas associated with a high risk of collision. By demonstrating that bees can form associations with environmental cues perceived both before and after collision events, we highlight the temporal flexibility in associative learning that contributes to their success in navigating complex natural environments. These findings also highlight the importance of negative reinforcement in shaping navigation behaviour. Our study demonstrates that - depending on the behavioural context - negative experiences are equally powerful drivers of adaptive navigation behaviour, revealing previously underappreciated cognitive flexibility in insect spatial learning.

## Acknowledgements

We would like to thank Jana Schäfer and Michelle Woestmann for their help during the data collection and the Deutsche Forschungsgemeinschaft (DFG) for funding the project.

## Data Availability Statement

The dataset and analysis pipeline for this study can be found in the repository “Learning to avoid collisions” https://gitlab.ub.uni-bielefeld.de/m.jeschke/learning_to_avoid_collisions

## References

Akpan, B. (2020). Classical and Operant Conditioning—Ivan Pavlov; Burrhus Skinner. Science Education in Theory and Practice, 71–84.

Avarguès-Weber, A., de Brito Sanchez, M. G., Giurfa, M., & Dyer, A. G. (2010). Aversive reinforcement improves visual discrimination learning in free-flying honeybees. PloS One, 5(10), e15370.

Avarguès-Weber, A., Deisig, N., & Giurfa, M. (2011). Visual cognition in social insects. Annual Review of Entomology, 56, 423–443.

Bertrand, O. J. N., Doussot, C., Siesenop, T., Ravi, S., & Egelhaaf, M. (2021). Visual and movement memories steer foraging bumblebees along habitual routes. The Journal of Experimental Biology, 224(11). 10.1242/JEB.237867

Bertrand, O. J. N., Lindemann, J. P., & Egelhaaf, M. (2015). A Bio-inspired Collision Avoidance Model Based on Spatial Information Derived from Motion Detectors Leads to Common Routes. PLoS Computational Biology, 11(11), 1–28.

Buatois, A., & Lihoreau, M. (2016). Evidence of trapline foraging in honeybees. The Journal of Experimental Biology, 219(16), 2426–2429.

Buehlmann, C., Dell-Cronin, S., Diyalagoda Pathirannahelage, A., Goulard, R., Webb, B., Niven, J. E., & Graham, P. (2023). Impact of central complex lesions on innate and learnt visual navigation in ants. Journal of Comparative Physiology. A, Neuroethology, Sensory, Neural, and Behavioral Physiology. 10.1007/s00359-023-01613-1

Burnett, N. P., Badger, M. A., & Combes, S. A. (2021). Wind and route choice affect performance of bees flying above versus within a cluttered obstacle field. In bioRxiv (p. 2021.10.08.463704). 10.1101/2021.10.08.463704

Chittka, L., Dyer, A. G., Bock, F., & Dornhaus, A. (2003). Psychophysics: bees trade off foraging speed for accuracy. Nature, 424(6947), 388.

Crall, J. D., Ravi, S., Mountcastle, A. M., & Combes, S. A. (2015). Bumblebee flight performance in cluttered environments: Effects of obstacle orientation, body size and acceleration. The Journal of Experimental Biology, 218(17), 2728–2737.

de Brito Sanchez, M. G., Serre, M., Avarguès-Weber, A., Dyer, A. G., & Giurfa, M. (2015). Learning context modulates aversive taste strength in honey bees. The Journal of Experimental Biology, 218(Pt 6), 949–959.

Eckel, S., Egelhaaf, M., & Doussot, C. (2023). Nest-associated scent marks help bumblebees localizing their nest in visually ambiguous situations. Frontiers in Behavioral Neuroscience, 17. 10.3389/fnbeh.2023.1155223

Egelhaaf, M. (2023). Optic flow based spatial vision in insects. Journal of Comparative Physiology. A, Neuroethology, Sensory, Neural, and Behavioral Physiology. 10.1007/s00359-022-01610-w

Fiala, A., & Kaun, K. R. (2024). What do the mushroom bodies do for the insect brain? Twenty-five years of progress. Learning & Memory (Cold Spring Harbor, N.Y.), 31(5), a053827.

Finke, V., Scheiner, R., Giurfa, M., & Avarguès-Weber, A. (2023). Individual consistency in the learning abilities of honey bees: cognitive specialization within sensory and reinforcement modalities. Animal Cognition, 26(3), 909–928.

Foster, D. J., & Cartar, R. V. (2011). What causes wing wear in foraging bumble bees? The Journal of Experimental Biology, 214(11), 1896–1901.

Giurfa, M., & Sandoz, J.-C. (2012). Invertebrate learning and memory: Fifty years of olfactory conditioning of the proboscis extension response in honeybees. Learning & Memory (Cold Spring Harbor, N.Y.), 19(2), 54–66.

Gonsek, A., Jeschke, M., Rönnau, S., & Bertrand, O. J. N. (2021). From Paths to Routes: A Method for Path Classification. Frontiers in Behavioral Neuroscience. 10.3389/fnbeh.2020.610560

Hussaini, S. A., Komischke, B., Menzel, R., & Lachnit, H. (2007). Forward and backward second-order Pavlovian conditioning in honeybees. Learning & Memory (Cold Spring Harbor, N.Y.), 14(10), 678–683.

Jeschke, M., Stahlsmeier, M., Egelhaaf, M., & Bertrand, O. J. N. (2025). Navigating in clutter: How bumblebees optimize flight behaviour through experience. The Journal of Experimental Biology. 10.1242/jeb.250514

Kamhi, J. F., & Barron, B. (2020). Vertical lobes mushroom bodies essential view-based navigation Australian Myrmeciaants. Current Biology, 30(17), 3432–3437.

Klein, S., Pasquaretta, C., Barron, A. B., Devaud, J.-M., & Lihoreau, M. (2017). Inter-individual variability in the foraging behaviour of traplining bumblebees. Scientific Reports, 7(1), 4561.

Knaden, M. (2020). Navigation: How the Recent Past Shapes Future Routes in Desert Ants. Current Biology: CB, 30(10), R435–R437.

Laska, M., Galizia, C. G., Giurfa, M., & Menzel, R. (1999). Olfactory discrimination ability and odor structure-activity relationships in honeybees. Chemical Senses, 24(4), 429–438.

Lehrer, M. (1993). Why do bees turn back and look? Journal of Comparative Physiology. A, Neuroethology, Sensory, Neural, and Behavioral Physiology, 172(5), 549–563.

Lehrer, M. (1994). Spatial vision in the honeybee: the use of different cues in different tasks. Vision Research, 34(18), 2363–2385.

Lihoreau, M., Chittka, L., Le Comber, S. C., & Raine, N. E. (2012). Bees do not use nearest-neighbour rules for optimization of multi-location routes. Biology Letters, 8(1), 13–16.

Lihoreau, M., Chittka, L., & Raine, N. E. (2010). Travel optimization by foraging bumblebees through readjustments of traplines after discovery of new feeding locations. The American Naturalist, 176(6), 744–757.

MaBouDi, H., Richter, J., Guiraud, M.-G., Roper, M., Marshall, J. A. R., & Chittka, L. (2025). Active vision of bees in a simple pattern discrimination task. eLife, 14. 10.7554/eLife.106332

Mountcastle, A. M., Alexander, T. M., Switzer, C. M., & Combes, S. A. (2016). Wing wear reduces bumblebee flight performance in a dynamic obstacle course. Biology Letters, 12(6). 10.1098/rsbl.2016.0294

Nityananda, V., & Chittka, L. (2015). Modality-specific attention in foraging bumblebees. Royal Society Open Science, 2(10), 150324.

Ravi, S., Bertrand, O., Siesenop, T., Manz, L. S., Doussot, C., Fisher, A., & Egelhaaf, M. (2019). Gap perception in bumblebees. The Journal of Experimental Biology, 222(2), 1–10.

Ravi, S., Siesenop, T., Bertrand, O., Li, L., Doussot, C., Fisher, A., Warren, W. H., & Egelhaaf, M. (2022). Bumblebees display characteristics of active vision during robust obstacle avoidance flight. The Journal of Experimental Biology. 10.1242/jeb.243021

Schwegmann, A., Lindemann, J. P., & Egelhaaf, M. (2014). Depth information in natural environments derived from optic flow by insect motion detection system: a model analysis. Frontiers in Computational Neuroscience, 8, 83.

Skorupski, P., & Chittka, L. (2010). Photoreceptor spectral sensitivity in the bumblebee, Bombus impatiens (Hymenoptera: Apidae). PloS One, 5(8), e12049.

Szyszka, P., Demmler, C., Oemisch, M., Sommer, L., Biergans, S., Birnbach, B., Silbering, A. F., & Giovanni Galizia, C. (2011). Mind the Gap: Olfactory Trace Conditioning in Honeybees. Journal of Neuroscience, 31(20), 7229–7239.

Webb, B. (2024). Beyond prediction error: 25 years modelingthe associations formed insect mushroom body. Learning & Memory, 31(5).

Woodgate, J. L., Makinson, J. C., Lim, K. S., Reynolds, A. M., & Chittka, L. (2016). Life-long radar tracking of bumblebees. PloS One, 11(8), 1–22.

Wystrach, A. (2023). Neurons from pre-motor areas to the Mushroom bodies can orchestrate latent visual learning in navigating insects. In bioRxiv (p. 2023.03. 09.531867). 10.1101/2023.03.09.531867

Wystrach, A., Buehlmann, C., Schwarz, S., Cheng, K., & Graham, P. (2020). Rapid Aversive and Memory Trace Learning during Route Navigation in Desert Ants. Current Biology: CB, 30(10), 1927–1933.e2.

