## Supplemental Figure 1 for "Learning to avoid collisions: Bees use cues perceived before and after a collision to prevent negative consequences during navigation in cluttered areas"

### Supplemental Material

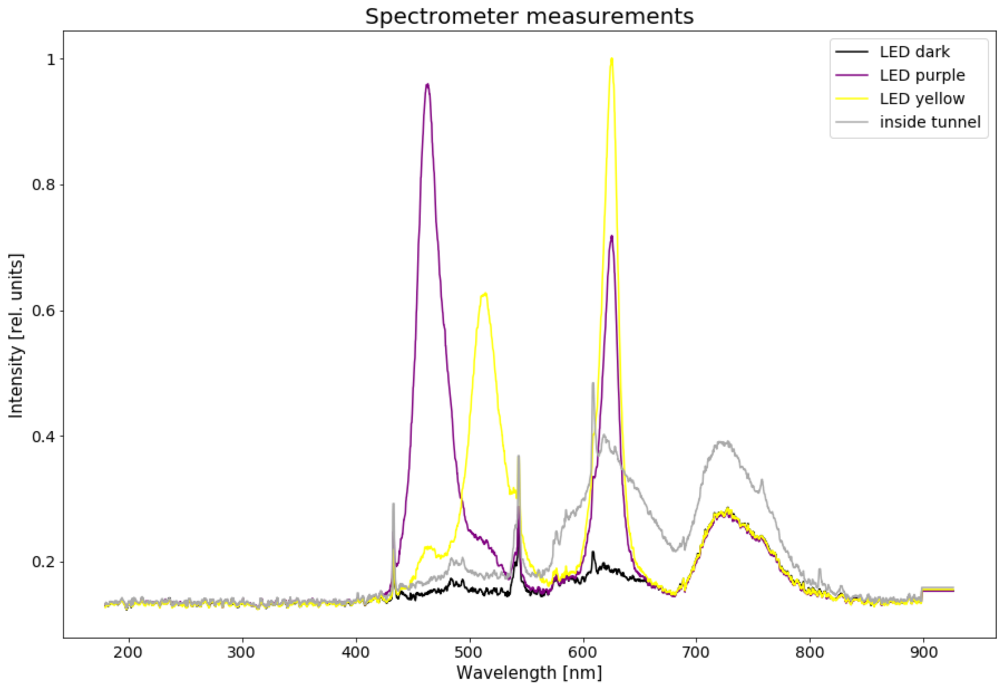


Supplement. Fig. 1: Spectrometer measurements for the colours displayed by the LED panel and the light inside the tunnel. Using the same light conditions as during the experiments, we measured the light spectrum of the purple LEDs (purple curve), of the yellow LEDs (yellow curve), the light when the LEDs are turned off (black curve) and the light inside the tunnel (grey curve).
